# A hyper-attenuated variant of Rift Valley fever virus (RVFV) generated by a mutagenic drug (favipiravir) unveils potential virulence markers

**DOI:** 10.1101/2020.10.16.342170

**Authors:** Belén Borrego, Alejandro Brun

## Abstract

In a previous work, we showed that favipiravir, a promising drug with antiviral activity against a number of RNA viruses, led to the extinction of RVFV from infected cell cultures. Nevertheless, certain drug concentrations allowed the recovery of a virus variant showing increased resistance to favipiravir. In this work, we characterized this novel resistant variant both at genomic and phenotypic level *in vitro* and *in vivo*. Interestingly, the resistant virus displayed reduced growth rates in insect cells and was highly attenuated but still immunogenic *in vivo*. Some amino acid substitutions were identified in the viral RNA-dependent RNA-polymerase (RdRp) gene and in the encoded IFN antagonist NSs gene, in catalytic core motifs and nuclear localization associated positions respectively. These data may help to characterize novel potential virulence markers, offering additional strategies for further safety improvements of RVF live attenuated vaccine candidates.

**SIGNIFICANCE STATEMENT:** Live attenuated virus vaccines usually provide long lasting immune responses upon administration. These vaccines are not recommended for use in immune compromised hosts, due to the presence of uncontrolled residual virulence. Cell culture virus propagation in the presence of mutagenic drugs often results in weakened virus lacking virulence as well as limited spreading capabilities. Here, we have characterized a mutagen-induced RVFV variant (40F-p8) that is not virulent in a highly sensitive mouse strain lacking antiviral response. The observed lack of virulence correlates with the presence of specific mutations along key residues in the viral genome, unveiling potential virulence determinants. Thus, 40F-p8 constitutes the basis for a novel RVFV vaccine strain with additional safety features.

## INTRODUCTION

Rift Valley fever virus (RVFV), a mosquito-borne bunyavirus belonging to the genus Phlebovirus in the *Phenuiviridae* family, causes an important disease in domesticated ruminants often transmitted to humans mainly through mosquito bites after epizootic outbreaks. Rift Valley fever is currently confined to the African continent and Southern parts of the Arabian Peninsula and Indian Ocean islands but its potential for spreading to other geographical areas, particularly linked to climatic change and globalization, has been widely remarked (1). In 2017, the World Health Organization ranked RVFV among the ten “most dangerous pathogens most likely to cause wide epidemics in the near future, requiring urgent attention” (http://www.who.int/blueprint/priority-diseases/en/). Currently, there is no available treatment or fully licensed RVF vaccines for use in non-endemic areas; consequently, developing of safer and effective control strategies intended also for human use is an active field of research.

The RVFV virion structure is formed by a lipidic envelope with two tightly packed membrane glycoproteins (Gn and Gc) arranged in an icosahedral lattice protecting an internal nucleocapsid composed by the viral nucleoprotein and a RNA dependent RNA polymerase (RdRp) bound to the viral RNA. The genome of RVFV is composed of three ssRNA segments of different size (<u>L</u>arge, <u>M</u>edium, <u>S</u>mall) with negative (L and M) or ambisense (S) polarity. In addition to the structural glycoproteins, the M segment encodes for two additional non-structural (NSm) proteins of 78kDa and 13-14kDa, while the S segment encodes the viral nucleoprotein and a 27kDa protein (NSs) considered the main virulence factor of the virus.

As many acute systemic viral infections, live-attenuated RVFV rapidly induce a long-lasting and broadly protective immunity after a single inoculation (2, 3). Therefore, vaccines based on attenuated virus remain as excellent candidates for a successful immunization program in the affected countries or as preventive control measure in countries with more elevated risk of disease introduction. For Rift Valley fever, live-attenuated vaccines have been generated either by random mutagenesis (4) or, more recently, by rationale deletion of virulence-associated genes using reverse genetics (5). In both cases, critical attenuating mutations or virulence determinants were identified, adding more available knowledge for further safety improvements.

However, the use of live attenuated vaccines in RVF endemic areas may still cause some safety concerns due to the possibility of genetic reassortment between closely related virus strains (6) due to the segmented genome of the virus. Although this phenomenon has been more often described for members of the orthobunyavirus genus (7) it is still considered as a potential drawback for live attenuated RVF vaccines. Additionally, although highly attenuated *in vivo*, these vaccines may retain some residual virulence, as shown upon experimental infection in immunocompromised lab animal models (8, 9) or in pregnant sheep when overdosed (10).

In a previous work, aimed to analyze the mutagenic effect of the nucleoside analog favipiravir on RVFV growth in *vitro*, we found that the propagation of the RVFV strain 56/74 in the presence of this drug led to virus extinction by a mechanism of lethal mutagenesis (11). Unexpectedly, at a dose of 40μM favipiravir, cytopathic effect (CPE) was detected in cell cultures after a lag phase (with no detectable CPE) of three consecutive blind passages. This finding was suggestive of incomplete or ineffective virus extinction leading to the selection of favipiravir-resistant variants. In this work, the virus recovered after eight serial passages in the presence of 40μM favipiravir (namely 40F-p8) was selected for further genomic, phenotypic and immunogenic characterization. The 40F-p8 virus displayed reduced growth rates in insect cells and, most interestingly, a “hyper attenuated” phenotype *in vivo*, as shown by the lack of virulence in the highly susceptible A129 mice (IFNAR^−/−^). These distinct features indicate that other virulence markers encoded in the RVFV genome remain to be characterized. Identification of these cryptic markers may help to strength the safety of live attenuated RVF vaccines.

## RESULTS

### 1. Phenotypic characterization of mutagen resistant RVFV 40F-p8 in cell culture

Firstly, we analyzed the kinetics and total virus yield of 40F-p8, the resistant virus recovered after 8 serial passages in Vero cells (ATCC-CL81) in the presence of 40μM favipiravir. The parental virus before (56/74) and after propagation along the same number of passages but in the absence of drug (56/74-p8) were included for comparison purposes. Since RVFV is an arbovirus, with mosquitoes playing an important role in the natural transmission cycle, infections were also carried out in *Ae.albopictus* clone C6/36 mosquito larvae derived cell line (ATCC-CRL1660).

Infections carried out in Vero cells showed growth curves similar for the three viruses (figure 1A). Titration of supernatants collected at different times post infection in several independent experiments showed only slight differences among the three viruses recovered. While the growth pattern of the virus passaged 8 times in the absence of drug showed no differences with the parental RVFV 56/74, the selected 40F-p8 virus displayed slightly faster growth, producing higher virus yields at 3-4 dpi that did not reach enough statistical significance (multiple t-test).

**Figure 1.**
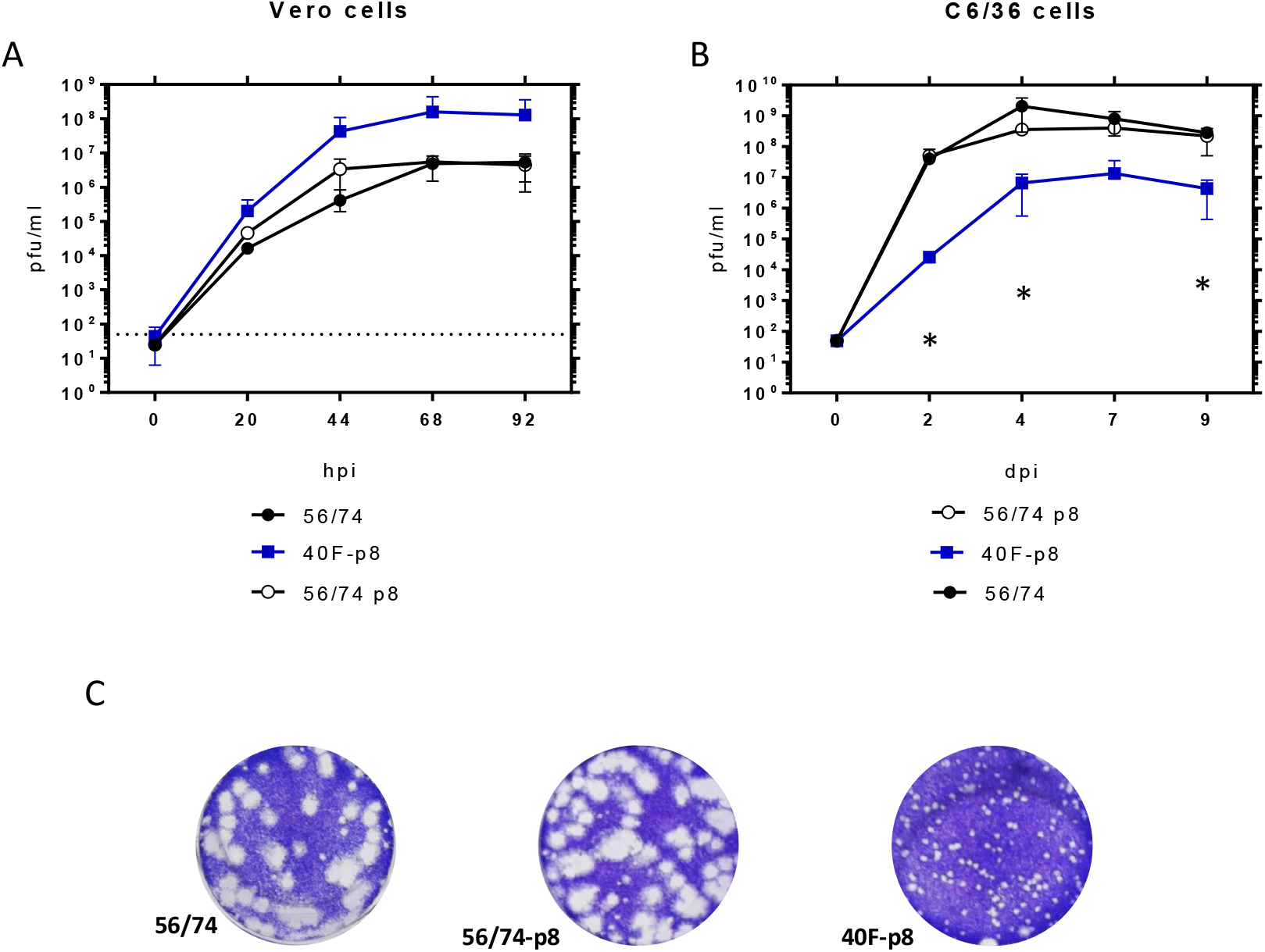
Kinetics of growth in mammalian and insect cells. Vero cells **(A)** and C6/36 mosquito cells **(B)** were infected at a MOI of 0.01 or 0.005 respectively. After one hour of adsorption the inoculum was removed, cells were washed and fresh medium was added. Supernatants were collected at different times post infection (pi) and titrated on Vero cell monolayers using a standard plaque assay using semisolid medium. Monolayers were fixed and stained 4 days post infection. Independent titrations were performed at least twice. Mean values ± SD are represented. Significance level was set to \**P* < 0.05 (multiple t-test using the Holm-Sidak method, with alpha=5.000%). Data shown correspond to one representative experiment. **C.** Plaque phenotypes of the indicated viruses on Vero cell monolayers. Infections were carried out in semisolid medium. Monolayers were fixed and stained 4 days post infection.

Conversely, both viral growth and final yield in C6/36 mosquito cells were clearly reduced for the selected 40F-p8 virus (figure 1B). Since infected mosquito C6/36 cells remain viable for longer times in cell culture than Vero cells, the analysis could be extended up to 9 days. In insect cells, the growth of 40F-p8 was significantly delayed, with viral titers of 10^4^ pfu/ml between 2-4 dpi, at least 3 log units lower than those rendered by the control viruses. Total virus yields at the latest points analyzed (7-9 days pi), although reaching a titer of 10^7^ pfu/ml, were still below the one reached by the parental RVFV 56/74 (>10^8^ pfu/ml). In contrast, no significant changes were found for 56/74-p8 with respect to the parental 56/74 virus.

Although the results obtained in Vero cells (fig 1A) did not show statistically significant differences among the three viruses, the plaque phenotype of 40F-p8 differed substantially, rendering smaller plaques than those produced by either the parental virus or by 56/74-p8 grown in the absence of favipiravir (figure 1C).

### 2. Analysis of the infectivity of RVFV 40F-p8 in A129 mice (IFNAR^−/−^)

Previous works have shown that viruses displaying some resistance to mutagenic antivirals are attenuated in vivo (12, 13). In fact, mutagen treatment has been often used as a procedure for virus attenuation. To test if the 40F-p8 virus was attenuated *in vivo* we performed an infection experiment using the interferon receptor deficient (IFNAR^−/−^) A129 strain of mice. Since these mice are unable to cope with an acute virus infection and are highly susceptible to RVFV infection (8, 14), we thought that they might offer a much more sensitive evaluation of the hypothesized attenuation of 40F-p8.

Groups of 5-6 mice were inoculated with different doses of each virus and monitored daily during 2 weeks for the development of signs of disease and survival (figure 2A). The virus propagated in the absence of drug (56/74-p8) caused 100 % of mortality 4 days after inoculation with 10^2^ pfu, while mice inoculated with the same dose of the parental RVFV 56/74 showed a survival rate of 40% (2/5). Although these data suggest a higher virulence than the parental strain 56/74, these results were not statistically significant (Mantel-Cox Log rank test) and were not further investigated. Higher doses of the parental strain 56/74 virus caused the death of the animals inoculated in the first 4 days after infection: 100% in those inoculated with 10^3^ pfu; 90% of those inoculated with 10^4^, with no survivors at day 10.

**Figure 2.**
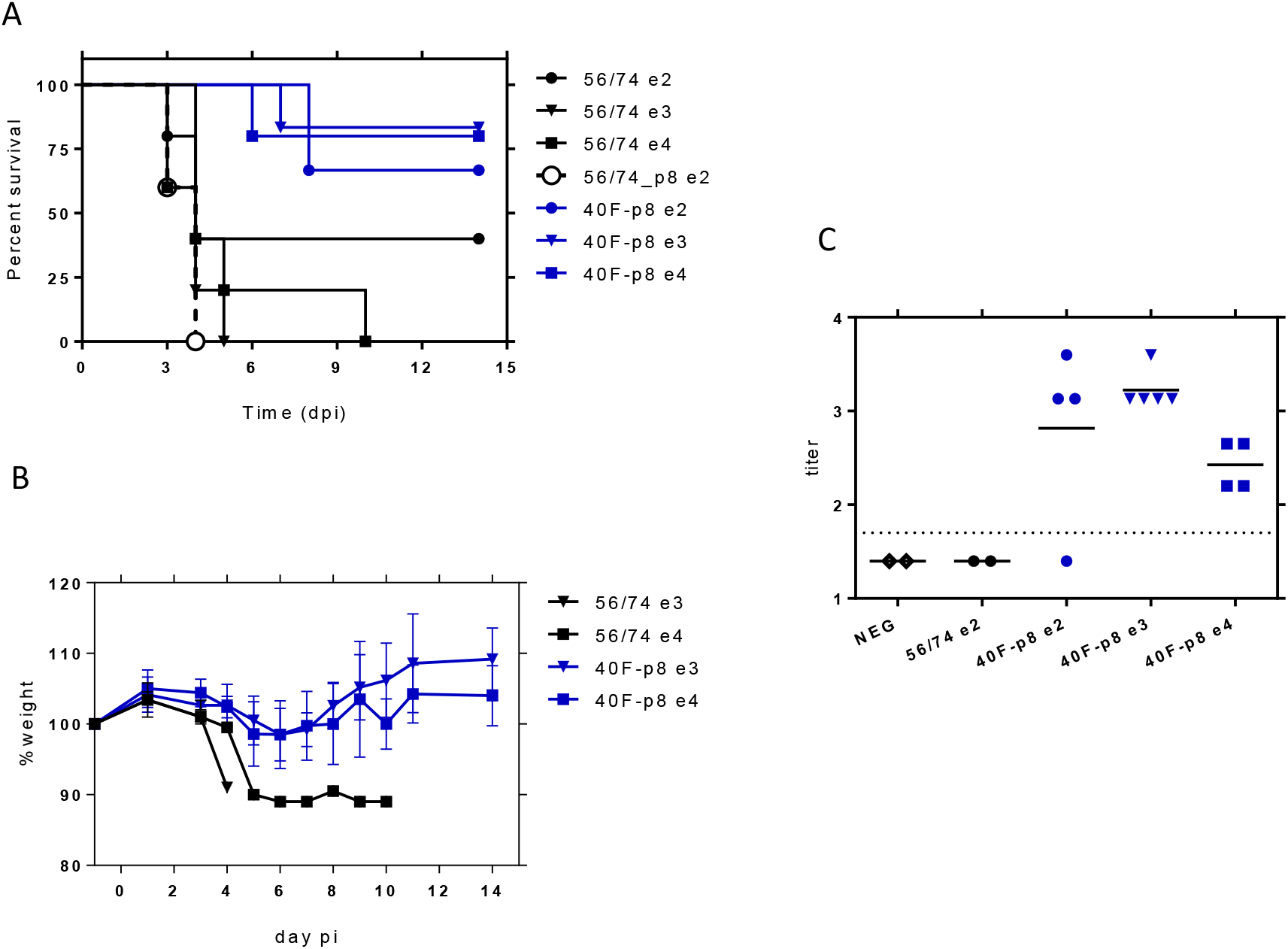
Analysis of the in vivo infectivity of 40F-p8 virus in A129 mice (IFNAR^−/−^). 5-6 month-old male mice (n=5 or 6) were inoculated IP with the indicated doses of virus. Animals were monitored daily during 14 days. **(A)** Survival rates. Curve comparison was performed using the Log-Rank (Mantel-Cox) test. **(B)** Body weight change in challenged mice with different doses. **C.** Detection of nucleoprotein specific antibodies by indirect ELISA. Titers are expressed as the last dilution of sera (log_10_) giving an OD_450_ reading over 1.0. Assay cut-off threshold was set to 1.7 (dotted line, corresponding to 1/50 serum dilution). Differences were assessed using a one-way ANOVA test.

Conversely, animals inoculated with 40F-p8 virus showed survival rates higher than 70% even at a high challenge dose (10^4^ pfu), with a significant number of survivors at the end of the experiment: 5/6 (83%) in those receiving 10^3^, and 4/5 (80%) in those inoculated with 10^4^. No signs of disease were observed in any of these animals except for a slight weight loss at days 3-5 pi (Fig 2B).

Serum samples collected at day 14 (end of the experiment) were tested by ELISA for the presence of anti-nucleoprotein N antibodies in the survivors as indicative of viral replication (FIG 2C). In some animals within groups receiving the lowest viral dose (10^2^ pfu) anti-N antibodies were undetectable, probably reflecting low or null levels of viral replication (2/2 in 56/74-inoculated mice; 1/4 in 40F-p8 inoculated mice). All animals inoculated with 10^3^ and 10^4^ pfu of 40F-p8, as well as three from the 10^2^ group developed specific anti-N antibodies.

Titers of anti-N antibodies did not show significant differences (ordinary one-way ANOVA) within the groups inoculated with 40F-p8, regardless of the dose received.

### 3. Immunogenicity and efficacy of 40F-p8 after RVFV challenge in 129 mice

The highly attenuated phenotype of the 40F-p8 virus displayed in immunodeficient A129 mice encouraged us to test its potential as a live attenuated vaccine in immune competent mice. With this aim, wild type 129SvEv mice were inoculated intraperitoneally (ip) with 10^4^ pfu of the 40F-p8 virus, and 4 weeks later they were challenged with a lethal dose (10^4^ pfu) of RVFV 56/74. After inoculation with 40F-p8 the mice did not show any sign of disease, not even significant weight variations (not shown). In serum samples collected 24 days after inoculation (pre-challenge samples), seven out of nine mice showed a strong neutralizing antibody response (FIG 3A). Anti-nucleoprotein N antibodies were detected in all these samples by indirect ELISA, including two samples that scored negative in the neutralization assay, although their anti-N antibody titers were slightly lower (FIG 3D, grey symbols). This suggested that the 40F-p8 virus replicated in all the inoculated mice at least to an extent enough to elicit an immune response. When subjected to a lethal challenge with the virulent strain 56/74, 100% of mice survived (*P* < 0.001, χ2 14.24, df 1) until the end of the experiment (FIG 3B, C) without apparent clinical display, including those in which neutralizing antibody titers had not been detected. In contrast, all mice in the control group became ill and died within day 4 (fig 3B, C).

**Figure 3.**
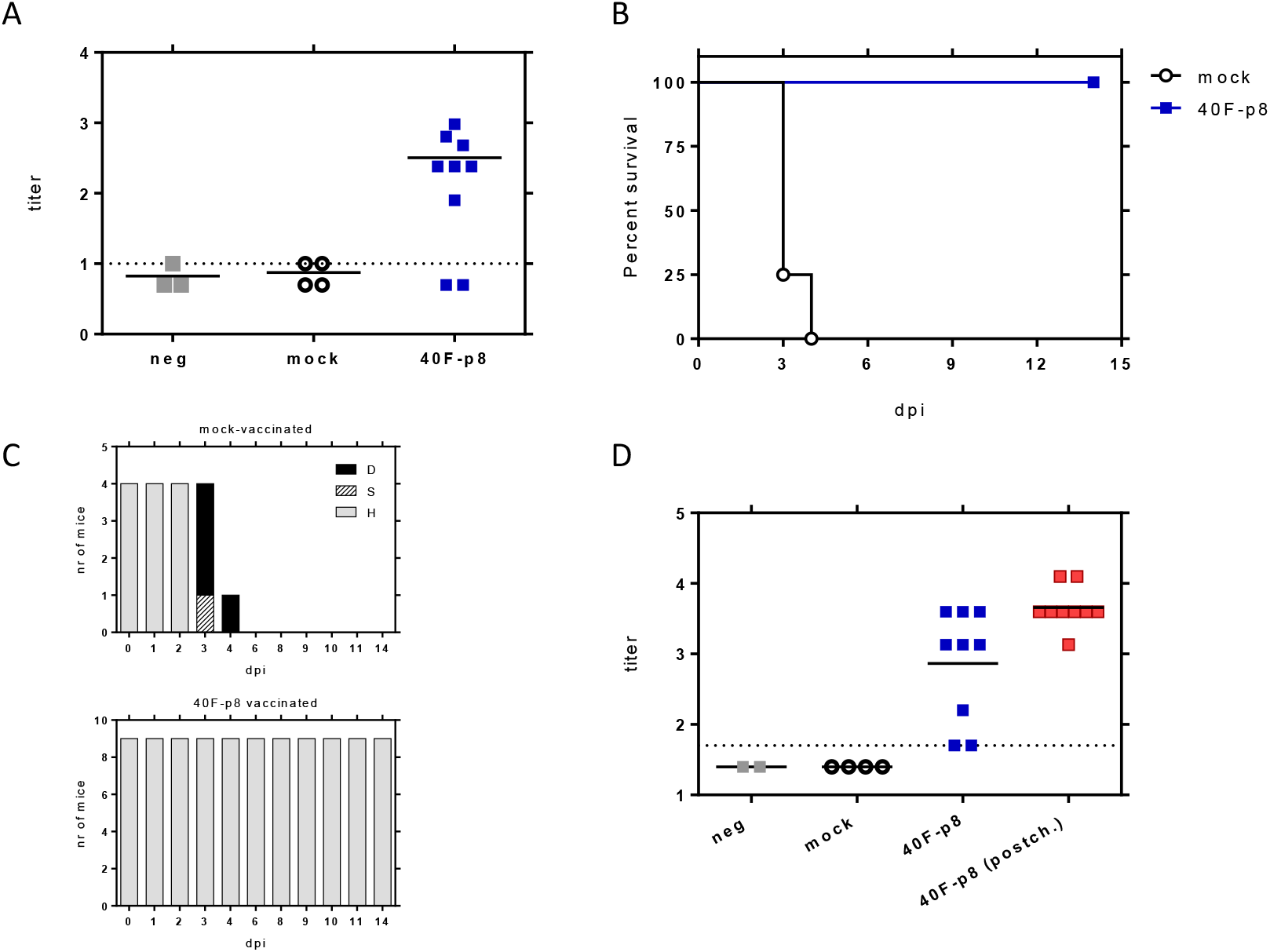
Analysis of the immunogenicity and efficacy upon lethal RVFV challenge of 40F-p8 virus. 11 month-old wild type 129Sv/Ev mice (n=9) were inoculated IP with 10^4^ pfu of 40F-p8. A group of mice was mock-inoculated (n=4). **A.** Microneutralization assay. Serum samples were taken at day 24 (pre-challenge samples) and titrated for neutralizing antibodies. Sera were assayed in 2-fold dilutions ranging from 1/10 to 1/1280. Titer was expressed as log_10_ of the highest serum dilution causing a CPE reduction of 50%. Assay cut-off was set to 1.0 (dotted line). Negative samples (rendering CPE in all wells at the first dilution assayed) were given an arbitrary value of 0.2. **B.** Survival of 129 mice upon RVFV challenge. Differences in survival assessed with the Log-Rank (Mantel-Cox) test. **C.** Morbidity upon challenge in vaccinated mice. The graph represents the clinical status of each mouse: D (dead): black bars; S (sick), hatched bars; H (healthy), grey bars. **D.** Detection of anti-nucleoprotein antibodies by ELISA. Each symbol corresponds to an individual animal, except for “neg” samples, corresponding to pools of preimmune sera. Black/blue symbols: pre-challenge samples. Red symbols: post-challenge samples (14 d pi). Sera were assayed in 3-fold dilutions ranging from 1/50-1/109350. Titers are expressed as the last dilution of sera (log_10_) giving an OD_450_ reading over 1.0. Assay cut-off threshold was set to 1.7 (dotted line, 1/50 serum dilution). For representation, negative samples were given an arbitrary value of 1.4.

Anti-N antibody titers were found increased upon virus challenge (FIG 3D, red symbols), suggesting a boosting effect in the primed mice. Altogether, these results show that in spite of its highly attenuated phenotype the 40F-p8 virus was able to replicate in immunocompetent 129Sv mice to levels allowing the induction of protective immune responses, even when no neutralizing antibodies are detected.

### 4. Genetic changes found in the selected viruses

In order to identify genetic changes that could be related to the observed phenotypic changes, the three RNA segments of the viral genome of the viruses obtained were sequenced, and deduced amino acid sequences were aligned and compared to that of the parental RVFV 56/74 strain. While no amino acid changes were found in the consensus sequence of the viral population recovered after eight passages without the drug, 47 nucleotide changes were found along the three RNA segments of 40F-p8 virus, leading to 24 amino acid substitutions (table I).

**Table I.**
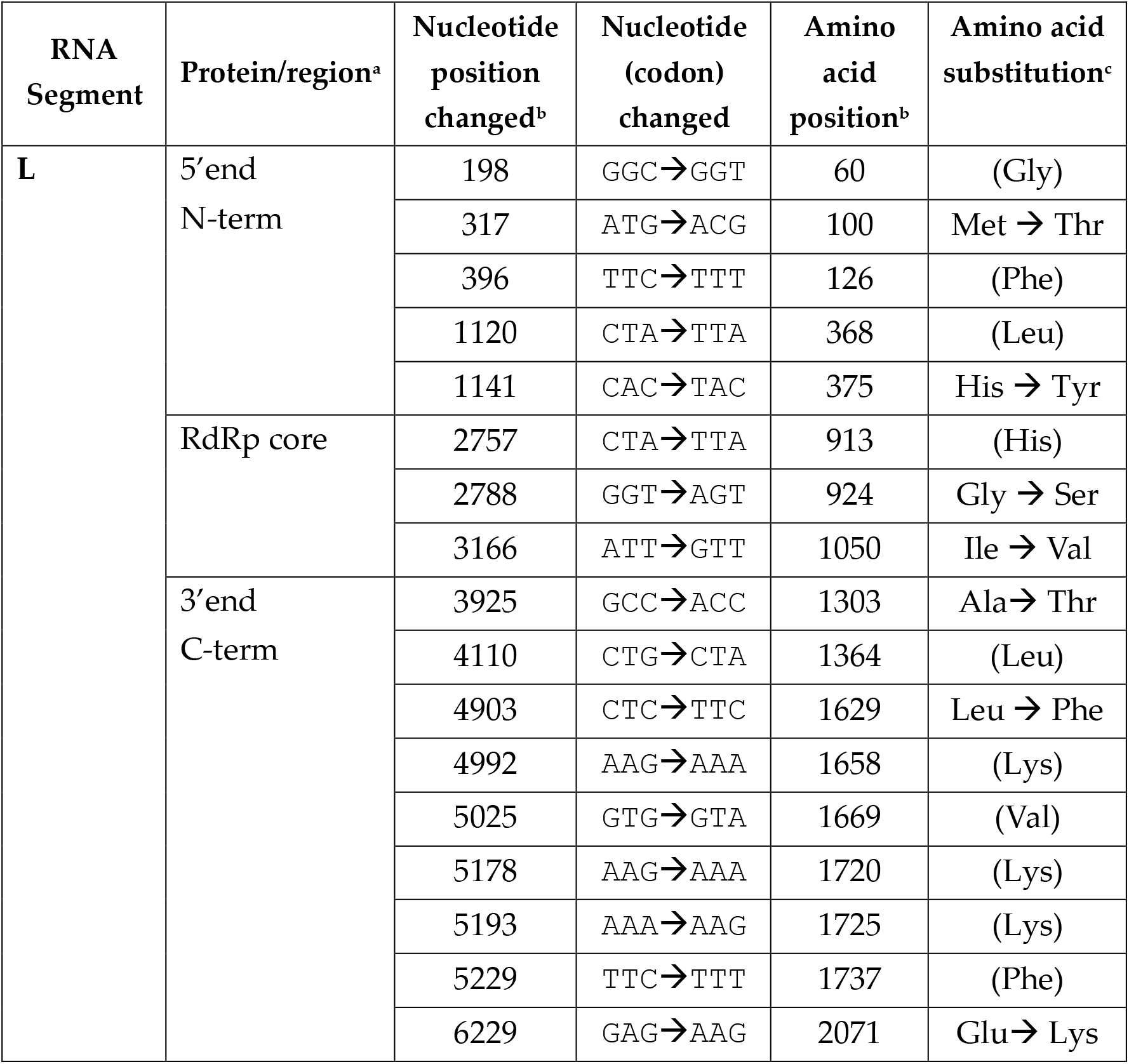
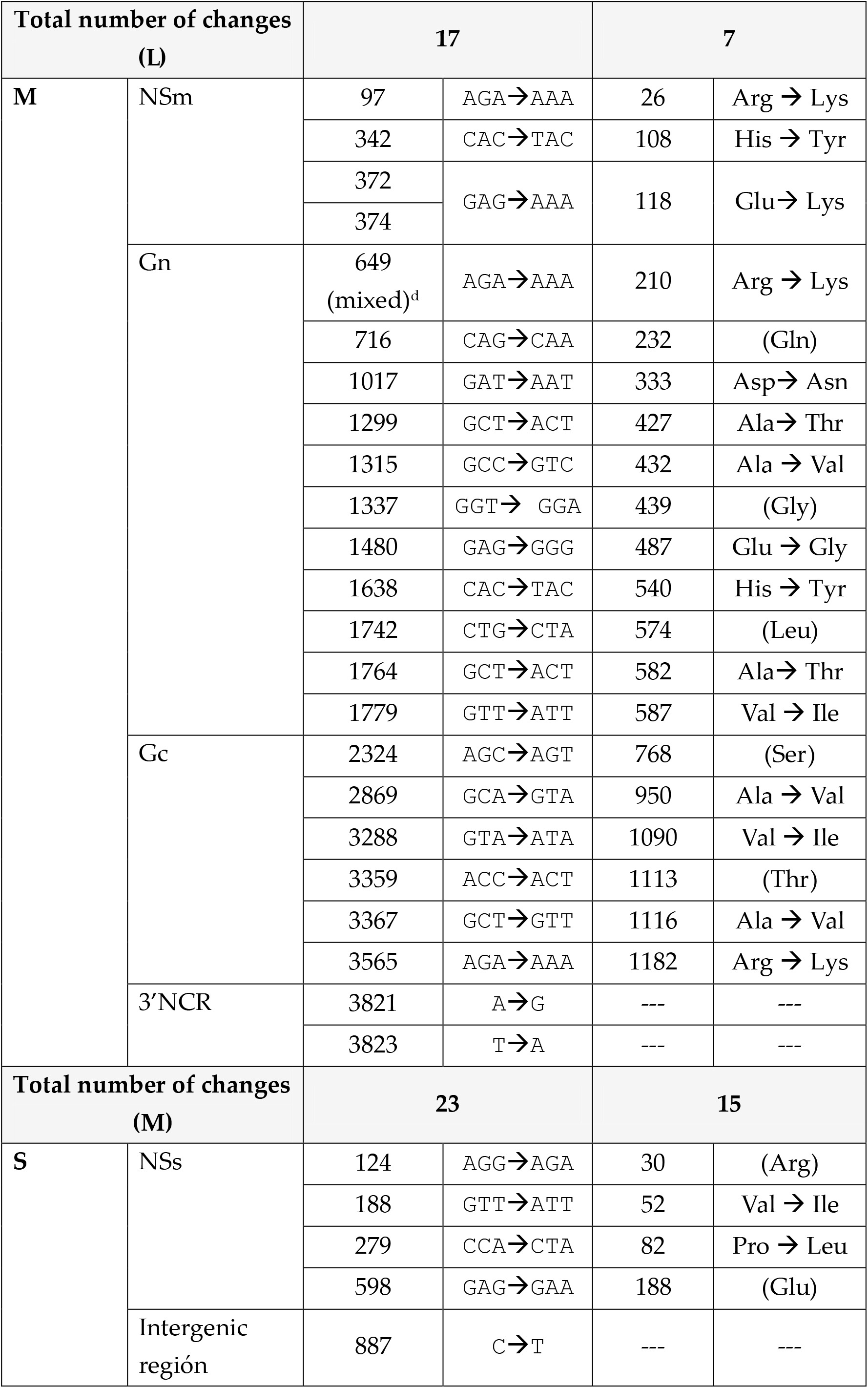
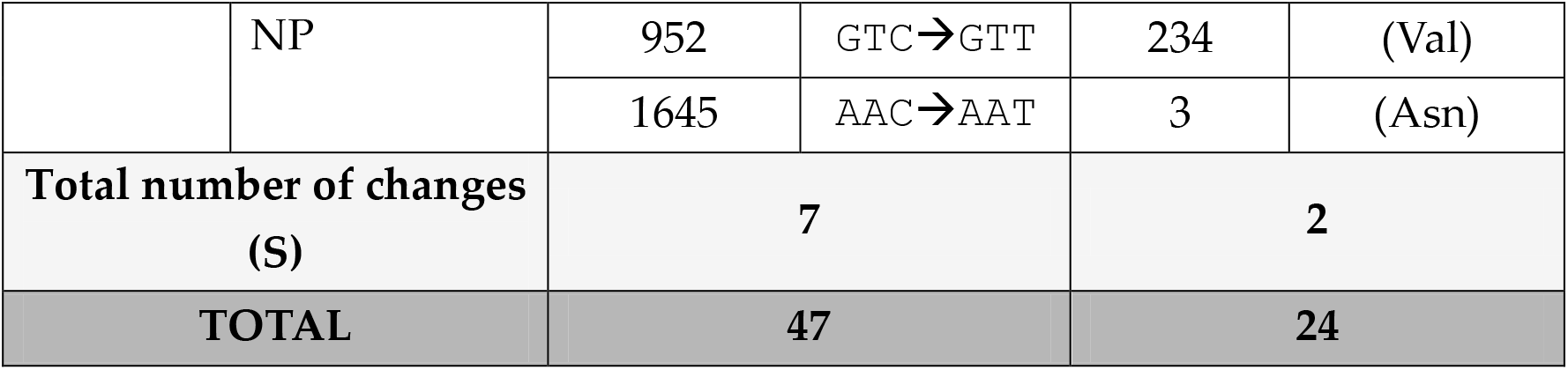
Total changes in the nt/aa sequence of 40F-p8 related to 56/74.

Most of the nucleotide changes were located in the ORFs of the corresponding RNA segments; only two changes were on 3’NCR of M segment and only one change was found in the intergenic region of segment S.

The six nucleotide changes found in the two S segment’ ORFs led to only 2 amino acid substitutions, both in the NSs protein: V52I and P82L. Interestingly P82 belongs to the second PXXP motif involved in the nuclear localization of the NSs protein and IFN-β activation (15). The nucleoprotein N was the only protein of 40F-p8 virus that showed an amino acid sequence identical to that of the parental virus, with only two (silent) nucleotide substitutions.

In the coding sequence corresponding to the M segment of 40F-p8 virus a total of 15 amino acid substitutions were identified, three in the NSm gene (R26K, H108Y, E118K), eight in the Gn coding sequence (R210K [mixed], D333N, A427T, A432V, E487G, H540Y, A582T, V587I) and four in the Gc ORF (A950V, V1090I, A1116V and R1182K). Interestingly, position R1182 (Gc) was previously involved in MP-12 virus attenuation (16).

The whole ORF of the L protein of the 40F-p8 virus showed seven amino acid substitutions, distributed along the entire sequence. Two changes were located in the N-term/third portion of the L-protein (M100T and H375Y); two were located in the C-term/third region (L1629F and E2071K), and the remaining three substitutions (G924S, I1050V and A1303T) corresponded to the central region of the protein. In particular, positions 924 and 1050 locate within the RpRd core (region 3 spanning amino acid positions 895-1206 as defined in (17), where conserved polymerase catalytic motifs A to H reside (18, 19).

Since the viral RNA polymerase is known to be a target of favipiravir, the drug used to select the 40F-p8 virus, we evaluated the level of conservation of the mutated residues that lay within the catalytic RpRd core, in an attempt to elucidate those involved in drug resistance. With this purpose we compared the L-protein sequences corresponding to 9 different RVFV strains corresponding to different genetic lineages (20) and also available sequences from 18 virus species belonging to the genus phlebovirus. Alignment ranged from amino acid position 895, the beginning of region 3 as described in (17), to position 1350, in order to cover also position 1303 (figure 4). The area around residue G924 (upper panel) was found to be highly conserved among all the sequences compared, as expected from its involvement on motif F (highlighted, consensus KQQHGGLREIYVMG). In particular, the residue G924 did not change in any of the sequences included. The area around position 1050 (central panel) showed a higher level of variation among sequences. In the RVFV isolates the residue 1050 was always isoleucine, while in the other phlebovirus species compared, other residues were found including valine (as displayed by 40F-p8). Finally, the region around A1303 displayed some degree of variation but this position was found to be extremely conserved in all the viruses included in the alignment.

**Figure 4.**
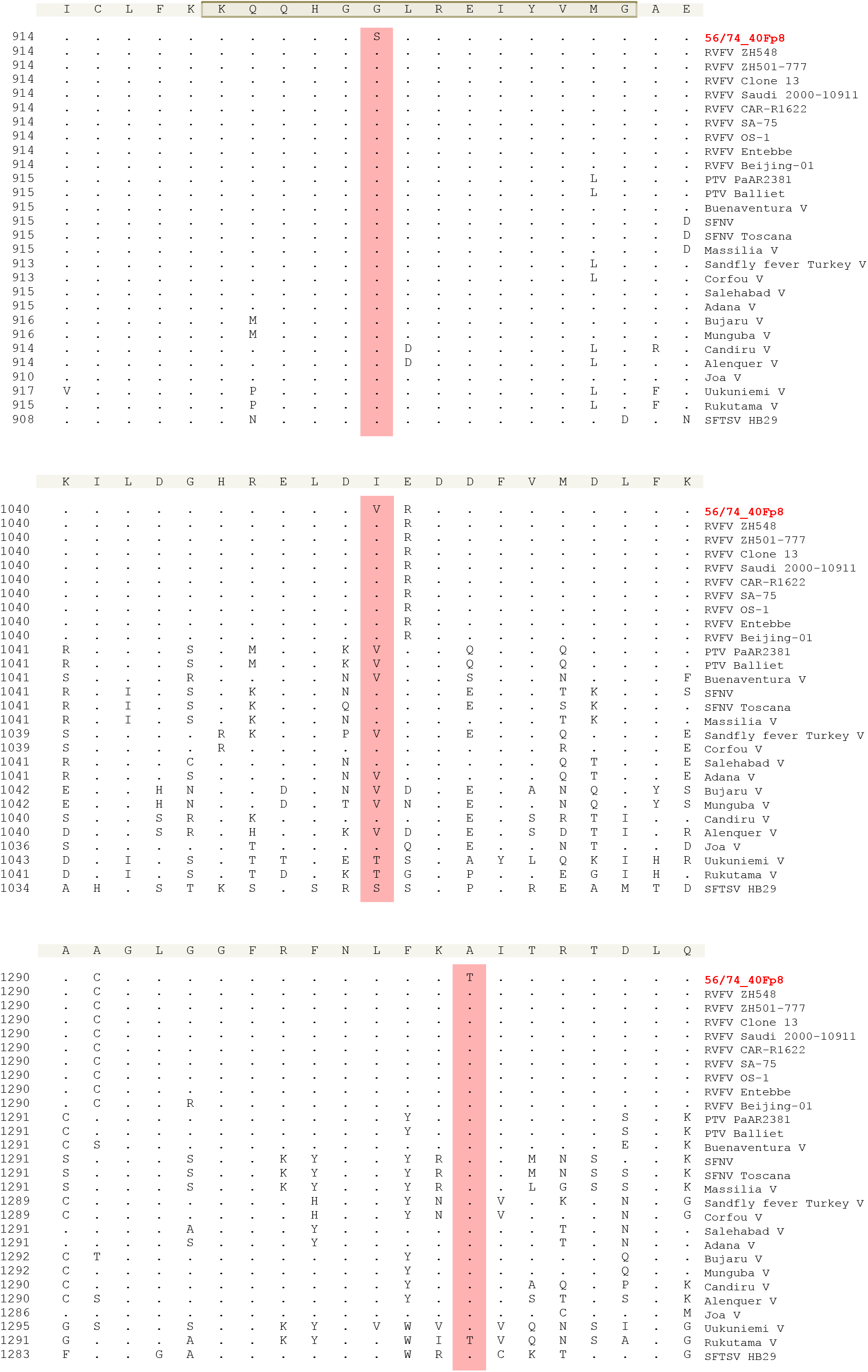
Multiple sequence alignment of phlebovirus species RdRps. Amino acid positions shown are 914-934 (upper panel), that contains the grey-shadowed F motif (19), 1040-1060 (central panel) and 1290-1310 (bottom panel). This numbering corresponds to the RVFVs sequences. En each panel, number in the left of each sequence indicates the amino acid position in the L protein for the corresponding virus. The alignment was generated with ClustalW using Laser gene software. Residues that match the sequence of the consensus (shadowed, top of each panel) exactly are hide as «-». Positions 924 (upper), 1050 (central) and 1303 (bottom) where 40F-p8 virus showed amino acid substitutions are highlighted. The sequences included and their database accession numbers are: RVFV ZH548 (DQ375403); RVFV ZH501 (DQ375408); RVFV Clone 13 (DQ375417); RVFV Saudi 2000-10911 (DQ375401); RVFV CAR-R1622 (DQ375423); RVFV SA-75 (DQ375428); RVFV OS-1 (DQ375398); RVFV Entebbe (DQ375429); RVFV Beijing-01 (KX611605); PTV PaAR2381 (KP272004); PTV Balliet (KR912212); Buenaventura V (KP272001); Sandfly fever Naples V(HM566172); SFNV-Toscana (NC_006319); Massilia V (EU725771.1); Sandfly Sicilian Turkey V (NC_015412.1); Corfou V (KR106177.1); Salehabad V (JX472403); Adana V (NC_029127); Bujaru V (KX611388); Munguba V (HM566164); Candiru V (NC_015374); Alenquer V (HM119401); Joa V (KX611391); Uukuniemi V (NC_005214); Rukutama V (KF892052); SFTS V HB29 (HM745930).

As shown in table I, some of the nucleotide changes found did not lead to an amino acid substitution in the corresponding protein (residues shown in parenthesis). Even though representing a small percentage out of the total codons, these 19 silent mutations were analyzed in terms of codon usage in different expression host organisms relevant for RVF infection (human, sheep, and mosquito) and in mice. To analyze whether the new codons present in the virus mutant 40F-p8 corresponded to a more or less represented codon usage, the frequencies per thousand of each mutated codon were compared with those of the parental virus 56/74. An unfavorable substitution was arbitrarily considered when ratios were ≤ 0.5 (i.e., the codon frequency in the mutant virus is half-represented in the corresponding organism related to the codon in the parental virus).

Based on this comparison (Supplementary table) we found that about half of the silent changes lead to unfavorable substitutions in both sheep and Aedes, while more similar in mice. If these silent nucleotide changes found in the 40F-p8 mutant virus exert some effect on gene expression of target organisms has not been further explored.

## DISCUSSION

Rift Valley fever is an emerging zoonotic disease relevant both for animal and human health. In Africa, RVF vaccines are available for livestock although different implementation policies are followed, depending on the epidemiological or socioeconomic situation of the countries (reviewed in (21). Veterinary vaccines in use are both classical inactivated vaccines as well as different live attenuated vaccines (LAVs). LAVs have proven to be more effective to control the disease but their use is still limited due to the risks associated to their residual virulence in pregnant sheep (10). In spite of vaccination campaigns, RVF outbreaks continue to occur, and it is accepted that most human RVF cases originate from infected animals. However, a licensed vaccine for human use in endemic countries is not yet available, not even for high risk populations exposed to RVFV contagion when handling infected animals during RVF outbreaks. A live-attenuated candidate vaccine, MP12, has been tested in clinical trials in humans, providing long-term protective immunity after a single dose, with low-to-moderate side effects. In 2013, the MP-12 vaccine received a conditional license for veterinary use in the U.S., but its application as human vaccine needs further improvement (reviewed in (14, 22). In Europe, no RVF vaccines, either for livestock or for human use, have been licensed.

The virus characterized in this work, 40F-p8, was generated in a similar manner to MP12, i.e., by serial passages in cell culture in the presence of a mutagenic agent (4, 11, 14); and as MP12, 40F-p8 displays a number of single mutations along the three RNA-genomic segments compared to the corresponding parental virulent strain. 40F-p8 showed an extremely high attenuation *in vivo*. When inoculated with a dose of 10^4^ pfu, the A129 (IFNAR^−/−^) mice showed a survival rate close to 80%. This was a quite remarkable finding, since this immune deficient strain of mice is highly susceptible to attenuated RVFV strains such as MP12, or even to NSm- or NSs - deletion based vaccines, with animals dying in a short period of time (8, 14). This attenuation however did not seem to impair its immunogenicity: when inoculated in immune competent mice, 40F-p8 was able to induce a protective immune response, even in the absence of detectable neutralizing antibodies in some of the animals. This fact underscores the potential of this virus as a candidate for the development of a safe live attenuated RVF vaccine and the interest of deciphering the changes leading to the observed hyper-attenuated phenotype. Similar data on the immunogenicity of 40F-p8 was confirmed in sheep (our unpublished observations) warranting further research to evaluate the protective efficacy of 40F-p8 virus as a vaccine for ruminants.

Even though expected because of the mutagenic effect of favipiravir (11, 23, 24), the high number of changes displayed by 40F-p8 (47 nucleotide changes, 24 amino acid substitutions) strongly hinders the identification of those responsible for the observed phenotype(s), especially attenuation. As previously reported for MP12, attenuation might be achieved by a combination of several individual amino acid changes (16). In fact, one of the many changes displayed by 40F-p8 in the glycoproteins (R1182K) affects the same residue already identified to contribute to attenuation of the strain MP12 (R1182G) (16). The other 14 aa substitutions in the M ORFs, although in positions not described (to our knowledge) to have a role in attenuation are to be investigated. Besides, silent nucleotide changes leading to misrepresented codons, even though representing a very low percentage, might have some effect on gene expression (25).

Changes in other proteins are more likely to be contributing to attenuation, for instance, those involving the NSs protein, known to be the main virulent factor, and in particular the P82L change, affecting the second PXXP motif of the protein (positions 82 to 85). Experiments in cells transiently transfected with mutant proteins where proline residues were substituted by alanine showed that the mutated protein did not reach the correct nuclear localization and lost their IFN-inhibiting activity (Billecocq 2004). However, our preliminary results indicate that the single P82L mutation does not impair nuclear NSs fibril-like formation (Borrego et al., unpublished data.)

Since RNA polymerases are targets of favipiravir, the mutagenic drug that led to selection of the attenuated 40F-p8 virus, changes found in this protein were especially interesting, in particular those in the central area corresponding to the RdRp core: G924S, I1050V and A1303T. In a structural model of L-protein, residue 924 is located within motif F in the RdRp core (17–19, 26). Motif F is involved in the binding of the incoming rNTP (27, 28) and plays a key role on the interaction of favipiravir with the viral polymerase, as already described for chikungunya virus (CHIKV), coxsackievirus B3 (CVB3), and influenza virus (29–31). The location of G924 within this motif strongly suggests that substitution G924S may be responsible for the partial resistance to favipiravir of 40F-p8 virus. Its high conservation in other phleboviruses supports its role as a key position on the RdRp. The nearby residues I1050V and A1303T may be compensatory or irrelevant changes, although the fact that A1303 is also highly conserved suggests that it also may play a relevant role.

The favipiravir resistance-phenotype could be actually contributing to attenuation. Viruses selected through resistance to mutagenic drugs may show attenuation in vivo because of the selection of high-fidelity polymerases (12, 13, 32). Higher fidelity polymerases give raise to viral populations with reduced genetic variation, thus decreasing their chances of adaptation to successful replication in different cell types, tissues or even hosts, a feature especially important for arboviruses whose life cycles involve both mammals and insects. Of note, one of the new features of 40F-p8 virus is the impaired growth displayed in mosquito-cultured cells, although whether this impairment occurs also *in vivo* needs to be determined. Because of the aforesaid, viruses with more reliable polymerases have been proposed as a novel strategy for the development of safer live attenuated vaccines, provided that immunogenicity is maintained (33–36). Furthermore, a vaccine virus with a higher fidelity polymerase would provide an additional safety measure by decreasing the chance of variation during its manufacturing or (35, 36) administered. If this is the case for the virus 40F-p8 is still to be determined. Work is in progress to elucidate the contribution of the individual changes, alone or in combination, to the phenotype(s) observed and their relationship with attenuation.

In summary, in this work we have characterized an RVFV variant, 40F-p8, selected by propagation in the presence of favipiravir. 40F-p8 displays a highly attenuated phenotype in IFNAR^(−/−)^ mice while retaining its immunogenicity, thus offering a promising RVF live attenuated vaccine candidate. 24 amino acid substitutions were found in the viral proteins, some of them in positions potentially involved in key processes of the viral cycle. The unequivocal identification of the changes responsible for attenuation as well as the other features observed for 40F-p8 should provide remarkable information on two important aspects for RVF control. Firstly, on the interaction of the favipiravir with the viral RdRp for a better understanding of the mechanisms of action of this antiviral drug and, secondly, on the unveiling of new in vivo markers of virulence that would open new strategies to improve the safety of RVFV live attenuated vaccines.

## MATERIALS AND METHODS

### Cells, viruses and infections

Vero cells (ATCC CCL-81) were grown in Dulbecco’s modified Eagle’s medium supplemented with 5%–10% fetal calf serum (FCS), and L-glutamine (2 mM), penicillin (100 U/ml) and streptomycin (100 μg/ml), in a humid atmosphere of 5% CO2 at 37°C. C6/36 *Aedes albopictus* cells (ATCC CRL 1660) were grown in Eagle’s Minimum Essential medium supplemented with 10% fetal calf serum (FCS), L-glutamine (2 mM), gentamicin (50 μg/ml), and MEM Vitamin Solution (Sigma) at 28°C. The origin of viruses used in this study has been described previously (11). Briefly, the South African RVFV strain 56/74 (parental virus) was serially passaged in the absence or presence of 40 μM favipiravir, and virus recovered after 8 passages in the presence of the drug was named as 40F-p8. Infections were performed as described (11).

### Animal experiments

Groups of 5-6 month-old transgenic 129Sv/Ev IFNAR^−/−^male mice (A129) or 11 month-old wild type 129Sv/Ev mice (B&K Universal) were inoculated intraperitoneally with different doses of the viruses, as indicated in the corresponding experiments. After viral inoculation, animals were monitored daily for weight and development of clinical signs, including ruffled fur, hunched posture, reduced activity and conjunctivitis. At the indicated time-points, animals were bled through the submandibular plexus. Sera were heat-inactivated at 56°C for 30 minutes and kept at −20°C until use. All mice were housed in a BSL-3 room with food and water supply *ad libitum*. All experimental procedures involving animals were performed in accordance with EU guidelines (directive 2010/63/EU), and protocols approved by the Animal Care and Biosafety Ethics’ Committees of INIA and Comunidad de Madrid (permit codes CEEA 2014/26, CBS 2017/15, PROEX 108/15 and PROEX192/17).

### Antibody assays

Neutralization assays were performed in 96-well culture plates following the OIE’s prescribed test for RVF (*OIE Terrestrial Manual 2012. Chapter 2.1.14*).

Briefly, sera were 2-fold diluted from 1/10 in DMEM containing 2% fetal bovine serum, mixed with an equal volume of infectious virus containing 100 TCID_50_ and incubated 30 minutes at 37°C. Then, a Vero cell suspension was added and plates were incubated for 4 days. Monolayers were then controlled for development of cytopathic effect (CPE), fixed and stained. Each sample was tested in 4 replica wells. Titer was expressed as the last dilution of serum causing CPE reduction in 50% of the wells.

For detection of antibodies against the nucleoprotein (N-protein), an in-house ELISA was performed. Briefly, ELISA plates were adsorbed with 100 ng/well of purified recombinant Trx-N protein produced in *E.coli* (37) and diluted in carbonate buffer (pH 9.6). After blocking with 5% skimmed-milk-PBS-0.05% Tween 20, sera were tested in duplicate in serial 3-fold dilutions starting at 1/50. Bound antibodies were detected with Goat Anti-mouse-IgG (H+L)-HRP Conjugated (BioRad) and bound conjugate was detected using TMB (Invitrogen/Life technologies) for 10 min, followed by one volume of stopping solution (3N H_2_SO_4_). Optical densities were measured at 450 nm (OD_450_).Titers are represented as the last serum dilution (log10) giving an OD ≥ 1.0.

### RNA extraction, RT-PCR, and nucleotide sequencing

RNA was extracted from the supernatants of infected cells using the Speedtools RNA virus extraction kit (Biotools B&M Labs) according to the manufacturer’s instructions. RT-PCR was performed using SuperScript IV Reverse Transcriptase (Invitrogen) and Phusion High-Fidelity DNA polymerase (Finnzymes), as directed by the manufacturers, using primers designed to amplify the S, M and L segments of the viral genome (Supplemental table S1). Overlapping PCR amplicons were purified and automatically Sanger-sequenced. For 3’- and 5’-ends of the RNA segments, a RACE approach was followed using the primers described in Supplemental table 3. Briefly, cDNAs from either genomic or antigenomic RNA ends were generated using Superscript IV enzyme mix. Upon RNAse H treatment, cDNAs were purified and subjected to A-tailing reaction using terminal deoxynucleotidyl transferase (TdT). After silica columm purification, PCR amplification with oligodT and RACE primers allowed sequencing of the genome ends. The Lasergene software suite (DNAstar) was used for analysis of the sequencing data.

## Statistical analysis

Data analysis was performed using GraphPad Prism software (version 6.0).

## ACKNOWLEDGEMENTS

We thank Francisco Mateos and Nuria de la Losa for excellent technical assistance. This work was supported by grants S2013/ABI-2906 (PLATESA), P2018/BAA-4370 (PLATESA2) from Comunidad de Madrid/FEDER and AGL2017-83326-R from Ministerio de Economía y Competitividad.

