## Supplemental tables for "A hyper-attenuated variant of Rift Valley fever virus (RVFV) generated by a mutagenic drug (favipiravir) unveils potential virulence markers"

**Supplementary data**

**Supplementary Table 1. Synonymous changes in 40F-p8 ORFs related to 56/74**

| | | **EXPRESSION HOST ORGANISMS** | | | |
|---|---|---|---|---|---|
| **RNA segment** | **residue position** | **MOUSE** | **HUMAN** | **SHEEP** | **AEDES alb.** |
| L | 60 | 0.53 | 0.47 | 0.49 | 1.16 |
| | 126 | 0.77 | 0.83 | 0.64 | 0.32 |
| | 368 | 0.81 | 1.04 | 1.45 | 0.41 |
| | 913 | 0.67 | 0.70 | 0.51 | 0.60 |
| | 1364 | 0.20 | 0.17 | 0.07 | 0.21 |
| | 1658 | 0.63 | 0.73 | 0.38 | 0.41 |
| | 1669 | 0.25 | 0.24 | 0.17 | 0.39 |
| | 1720 | 0.63 | 0.73 | 0.38 | 0.41 |
| | 1725 | 1.58 | 1.40 | 2.63 | 2.44 |
| | 1737 | 0.77 | 0.83 | 0.64 | 0.32 |
| M | 232(Gn) | 0.34 | 0.34 | 0.40 | 0.43 |
| | 439(Gn) | 1.47 | 1.51 | 1.03 | 0.98 |
| | 574(Gn) | 0.20 | 0.17 | 0.07 | 0.21 |
| | 768(Gc) | 0.63 | 0.61 | 0.39 | 0.73 |
| | 1113(Gc) | 0.71 | 0.67 | 0.25 | 0.36 |
| S | 30(NSs) | 0.97 | 1.01 | 0.78 | 1.40 |
| | 188(NSs) | 0.67 | 0.71 | 0.48 | 1.24 |
| | 234(N) | 0.68 | 0.75 | 0.69 | 0.68 |
| | 3(N) | 0.75 | 0.86 | 0.33 | 0.47 |

Ratio between genetic codon frequencies in the indicated expression host organisms. Ratio between frequencies per thousand corresponding to codon present in the virus mutant and codon present in the parental virus, according to https://www.genscript.com/tools/codon-frequency-table, https://www.kazusa.or.jp/codon/. For ratios **below 0.5** the nucleotide change is considered unfavorable in terms of codon usage (shadow).

**Supplementary Table 2. List of primers used in this study (RT-PCR and/or sequencing).**

| Oligo name | sequence (5′-3′) | nt positions[a] (L segment) | orientation |
|---|---|---|---|
| L seg 5′ end | ACACAAAGGCGCCCAATCATGGATTCTATA | 1-30 | antigenomic |
| 716fwd | CAGTCACAGAATCTGTG | 716-732 | antigenomic |
| L-fwd 1028ag | ATGGCAAAGATCTGTGC | 1028-1044 | antigenomic |
| L-rev 2300g | AAGTAACCATTGTAACAACA | 2300-2281 | genomic |
| RdRp central-fwd | GCTATCCAGAAGTTTGAGGATTG | 2701-2723 | antigenomic |
| L-seg-fwd[b,d] | TTCTTTGCTTCTGATACCCTCTG | 2872–2894 | antigenomic |
| L-seg-rev[b,d] | GTTCCACTTCCTTGCATCATCTG | 3006-2984 | genomic |
| RdRp central-rev | CTGATTTGCAGAAGCTATATGCT | 3960-3938 | genomic |
| 3817 fwd | GAGTATAAGAAGGCAGT | 3817-3833 | antigenomic |
| 4553 fwd | TCACTATCTTTAATGAG | 4553-4569 | antigenomic |
| 5455 fwd | CCATTTGGATGCCCAGTTTATAT | 5455-5477 | antigenomic |
| 5583 rev | CTGATGAGAATGGTGCT | 5583-5567 | genomic |
| Q3′25nts | TTGTAGCACTATGCTAGTATCAGAA | 6385-6361 | genomic |

| Oligo name | sequence (5′-3′) | nt positions[a] | orientation |
|---|---|---|---|

| | | (M segment) | |
|---|---|---|---|
| (-2)RTsm1 | GTTTTATTAACAATTCTAACCTCGGTT | 27-53 | antigenomic |
| MRV1ag[c] | CAAATGACTACCAGTCAGC | 772-790 | antigenomic |
| RTsm2 | CAAGTGAACAGGGAAATAGGATGG | 1953-1976 | antigenomic |
| sm2 | GAACATGCTGATGCATATGAGACA | 2095-2072 | genomic |
| sm3 | GACCCATCCTTATTGTGGGCAGAG | 3223-3200 | genomic |
| sm4 | ATCTGGCAAAAGACGATTACGC | 3838-3817 | genomic |
| EM-rev[d] | CACAAAGACCGGTGCAAC | 3884-3867 | genomic |
| EM-fwd[d] | GAATCAACAGTTGTGAATCC | 3405-3424 | antigenomic |

| Oligo name | sequence (5′-3′) | nt positions[a] (S segment) | orientation |
|---|---|---|---|
| NS0g | ACACAAAGACCCCCTAGTG | 1-19 | genomic |
| NS2g[e] | GATTTGCAGAGTGGTCGTC | 62-80 | genomic |
| revS[d] | CGGACTTGGAGACTTTGCATCA | 241-262 | genomic |
| fwdS[d] | TTCTTTCAGATTGGGGAACCTTGT | 36-338 | antigenomic |
| NSca[e] | CCTTAACCTCTAATCAAC | 841-824 | antigenomic |
| ss1 | AGCCACTTAGGCTGCTGTCTTGT | 908-930 | genomic |
| RTss1 | CTATTACAATAATGGACAACTATCAAGAGC | 1662-1633 | antigenomic |
| NP0ag | ACACAAAGCTCCCTAGAGATA | 1689-1669 | antigenomic |

[a]Nucleotide positions according to RVFV MP12 sequence (GB accession: DQ380208.1) [b] Busquets et al. Vector Borne Zoonotic Dis. 2010 10(7):689-96.[c]Takehara et al Virology. 1989 169(2):452-7; [d]Mwaengo et al. Virus Res. 2012 169(1):137-43; [e]Sall et al. Clin Diagn Lab Immunol. 2002 9(3):713-5

### Supplementary table 3. List of primers used for RACE

| Oligo name | sequence 5′-3′ | Positions[a] (segment) | orientation |
|---|---|---|---|

| | | | |
|---|---|---|---|
| Lseg cRNA | AGTTTTGATAATCATGTC | 366-349 | genomic |
| Lseg vRNA | CTTTTGATACATCTGA | 6011-6026 | antigenomic |
| Mseg cRNA | GTTTCTCTCCCTATTTG | 409-393 | genomic |
| Mseg vRNA | TCCTCCTTATATATCTTG | 3544-3561 | antigenomic |
| Sseg cRNA | TATTAGATCAATAAGTCT | 313-296 | antigenomic |
| Sseg vRNA | TCTTTCGACATTTTCAT | 1419-1435 | genomic |

[a] nt positions according to the strain 56/74 sequence elucidated in this work (GB annotation pending)
